# Zinc Starvation Drives Respiratory Remodeling and Metabolic Adaptations Associated with Cystic Fibrosis in *Pseudomonas aeruginosa*

**DOI:** 10.64898/2026.07.15.738699

**Authors:** Emma Michetti, Valerio Secli, Francesca Pacello, Federico Iacovelli, Marta Mellini, Chiara Scribani Rossi, Serena Rinaldo, Francesca Cutruzzolà, Giordano Rampioni, Serena Ammendola, Andrea Battistoni

## Abstract

Host nutritional immunity restricts microbial growth by altering metal availability. In patients with cystic fibrosis (CF), microorganisms inhabiting the thick airway mucus experience severe nutrient limitation, particularly zinc (Zn) restriction, which plays a critical role in limiting lung colonization. However, *Pseudomonas aeruginosa,* a major contributor to morbidity and mortality in CF, employs multiple strategies to overcome Zn deficiency and persist within the airways. Understanding how *P. aeruginosa* adapts to Zn deprivation may facilitate the development of antimicrobial approaches targeting Zn homeostasis. In this study, we characterized the physiological and transcriptional adaptations that support *P. aeruginosa* survival under Zn-limited conditions. Transcriptomic analysis of a *znuAzrmB* mutant unable to efficiently acquire Zn revealed widespread repression of pathways involved in central carbon metabolism, motility and virulence. Notably, Zn limitation promoted extensive respiratory remodeling, characterized by a shift toward anaerobic metabolism, induction of denitrification pathways, altered terminal oxidase expression, and reduced oxygen consumption. These metabolic changes correlated with decreased ATP production, altered membrane potential, and increased aminoglycoside tolerance. Furthermore, the Zn-starved mutant exhibited reduced production of quorum-sensing molecules, redox imbalance and altered oxidative stress responses. Many of these adaptations resemble those observed in *P. aeruginosa* isolated from CF sputum, suggesting convergence towards a common host-adapted physiological state. Collectively, these findings identify Zn starvation as a major driver of bacterial physiological remodeling in CF conditions and reveal a previously unrecognized link between Zn limitation, respiratory reprogramming, and the emergence of persistence-associated traits in *P. aeruginosa*.

**Importance:** During infection, the host restricts Zn availability as part of nutritional immunity, but how this influences the physiology of *Pseudomonas aeruginosa* remains poorly understood. Here we show that severe Zn limitation triggers a coordinated metabolic program that extends far beyond Zn acquisition, encompassing respiratory remodeling, altered energy metabolism, redox imbalance, and increased tolerance to aminoglycosides. Remarkably, many of these changes resemble transcriptional and physiological traits previously described in bacteria adapted to cystic fibrosis airways, identifying Zn limitation as a key environmental signal that contributes to chronic infection–associated phenotypes. These findings broaden our understanding of how metal availability shapes bacterial physiology and suggest that targeting Zn homeostasis may influence both bacterial persistence and antibiotic susceptibility.

## Introduction

Successful persistence of bacterial pathogens within host tissues depends on their ability to remodel cellular physiology in response to environmental constraints. During chronic infections, many pathogens shift from actively growing states toward specialized physiological programs characterized by altered metabolic activity. This adaptive strategy is particularly evident in chronic airway infections caused by *Pseudomonas aeruginosa*, an opportunistic pathogen that significantly contributes to mortality in cystic fibrosis (CF) patients (1).

In the CF lung, *P. aeruginosa* experiences a complex and heterogeneous environment shaped by nutrient limitation, oxidative stress, fluctuating oxygen availability, and antimicrobial pressures imposed by both host immunity and antibiotic therapy (2,3). Analyses of clinical isolates and sputum-derived populations have revealed extensive remodeling of central metabolism, respiratory pathways, motility, and virulence-associated functions, reflecting adaptation to conditions that favor long-term persistence (4,5). While some of these phenotypes have been linked to nutrient availability, the environmental signals that initiate and coordinate these adaptive responses remain incompletely understood.

One of the most common host-imposed stresses encountered during infection is the restriction of trace metals through nutritional immunity, an innate immune defense mechanism highly conserved across evolutionarily diverse organisms (6–8). Zinc (Zn) is of particular importance because it is required for many bacterial cellular processes, including enzyme function, transcriptional regulation, and virulence (9–11). Elevated levels of the metal-chelating protein calprotectin, released by infiltrating neutrophils, contribute to Zn sequestration in the CF airways and generate localized environments in which Zn availability becomes limiting for bacterial pathogens (12,13). *P. aeruginosa* maintains Zn homeostasis through multiple highly efficient uptake systems, controlled by the master regulator Zur, including the high-affinity ZnuABC transporter, the ZrmABCD pseudopaline system, and additional components involved in Zn acquisition and Zn-sparing adaptation (9,14,15). Consistently, transcriptomic analyses of *P. aeruginosa* recovered from CF sputum have repeatedly identified a strong induction of the Zur regulon, indicating that Zn limitation is a physiologically relevant stress encountered during chronic CF lung infection (4,16,17). The importance of Zn acquisition for bacterial fitness within the host has prompted the development of therapeutic strategies that exploit Zn uptake pathways for antimicrobial delivery (18,19).

Although substantial progress has been made in elucidating the molecular mechanisms underlying Zn homeostasis and the bacterial response to Zn deprivation, it remains unclear to what extent Zn limitation contributes to the large-scale physiological remodeling associated with chronic CF lung infection. In this study, we investigated the global physiological adaptation of *P. aeruginosa* to severe Zn starvation using a mutant lacking the two major Zn uptake systems, *i.e.*, ZnuABC transporter and the ZrmABCD pseudopaline system. Integrating transcriptomic analyses and physiological profiling, we show that Zn limitation triggers extensive cellular reprogramming affecting central metabolism, respiratory function, redox homeostasis, quorum sensing, and antibiotic susceptibility. Together, these findings reveal how Zn limitation drives a coordinated physiological program that recapitulates key features of *P. aeruginosa* adaptation to the CF lung environment.

## Results

### Transcriptional adaptations of *P. aeruginosa* to severe Zn deficiency

To systematically characterize the transcriptional changes that support *P. aeruginosa* adaptation under Zn-limited conditions, we performed a comparative RNA-seq analysis of the PA14 wild-type strain and the *znuAzrmB* mutant grown under Zn-depleted conditions (**Table S1**). The analysis revealed 600 genes that were differentially expressed in the *znuAzrmB* mutant compared to the wild-type strain with Log_2_ FC ≥ |1.3| and an adjusted p-value ≤ 0.05 (**Fig. 1, panel A**). Functional enrichment analysis was performed using Clusters of Orthologous Genes (COGs), Kyoto Encyclopedia of Genes and Genomes (KEGG) pathways, and manually curated gene classes (**Fig. 1, panel B**).

**Fig. 1:**
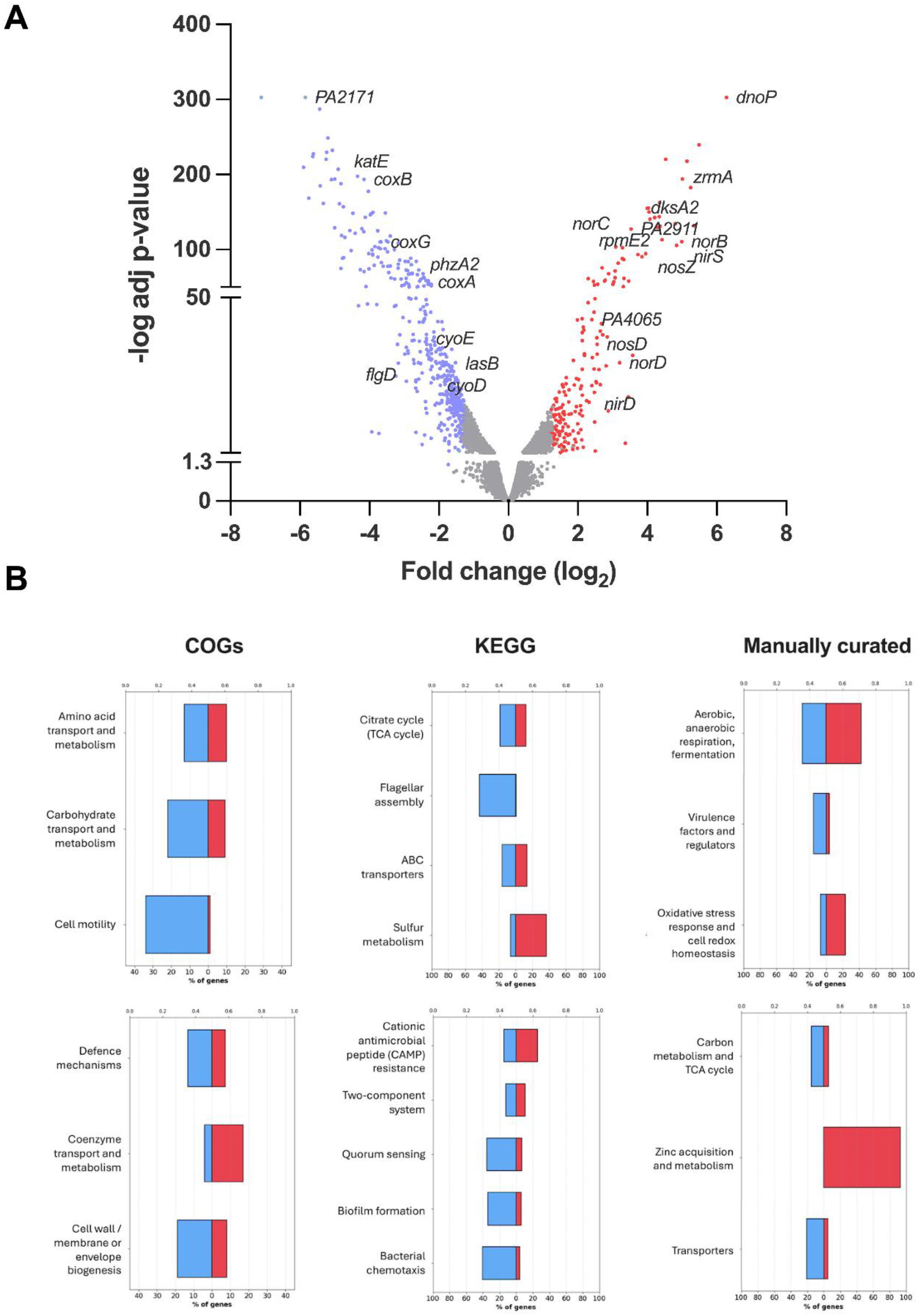
Transcriptional modulation of the Zn-starved *P. aeruginosa*. (**A**) Volcano plot of the differentially expressed genes of the *znuAzrmB* strain compared to PA14 wild type, grown in E-VBMM. Red and blue dots indicate significantly upregulated and downregulated genes, respectively, with log_2_ FC < |1.3| and adjusted p-value > 0.05. Grey dots represent genes not significantly regulated. (**B**) Gene enrichment analyses based on COGs classification systems, KEGG pathways, and manually curated gene classes. Gene association for each category was obtained from the *Pseudomonas.com* database. In each plot, the percentage of genes significantly upregulated (red bars) or downregulated (blue bars) associated with each functional category is reported, with an adjusted p-value ≤ 0.05 (Bonferroni correction for multiple testing).

As expected, Zur-regulated genes involved in Zn import and adaptation to Zn starvation were strongly upregulated in the *znuAzrmB* mutant. In contrast, downregulated genes were significantly enriched in pathways related to carbohydrate transport and metabolism. These included *PA14_22980–PA14_23000*, as well as *oprB* and *gapA*/*gapB*, involved in glucose uptake and metabolism; *kguT*, *kguK*, *PA14_34630, gntR* and *gntK*, associated with gluconate transport and metabolism; *glgABP*, encoding enzymes involved in glycogen biosynthesis and degradation; and the *PA14_36740–PA14_36730* locus, implicated in the biosynthesis of the osmoprotectant trehalose. In addition, genes associated with pyruvate metabolism were downregulated, including *pdhB* and *PA14_19900*, which putatively encode components of the pyruvate dehydrogenase complex. Consistent with these changes, a broader repression of central carbon metabolism was observed, including the tricarboxylic acid (TCA) cycle gene *acnA*. Genes involved in amino acid metabolism were also differentially regulated in the *znuAzrmB* mutant, with induction of methionine and serine biosynthesis genes, including *metE* and *serA*, respectively, whereas genes of the glycine cleavage pathway (*gcvT2* and *gcvP1*) were downregulated. Furthermore, the Zn-starved mutant exhibited reduced expression of genes associated with multiple virulence determinants, including motility and chemotaxis genes (*cheA*, *cheB*, *cheW*, and flagellar genes *flgBCDE*), quorum sensing (QS)-regulated virulence determinants (*lasA*, *lasB*, *aprA*, and phenazine biosynthesis operons *phzA1-G1* and *phzA2-G2*, as well as *phzH*, *phzS*, and *phzM*), biofilm-associated genes (*cdrA* and *lecB*), and multidrug efflux systems, including *mexAB*-*oprM* and the *PA14_40230*–*PA14_40260* operon (*opmL*). In contrast, genes associated with cationic antimicrobial peptide (CAMP) resistance were significantly upregulated, including the operon *PA14_63110*–*PA14_63160,* which includes the PmrAB two-component system regulating LPS modifications and stress response (20). Moreover, the transcriptional profile of the Zn-starved mutant revealed induction of sulfur acquisition and metabolism genes, including *ssuABC*, *ssuD* and *tauD,* which are required for the utilization of taurine and alkanesulfonates as sulfur sources. Genes involved in oxidative stress responses and redox homeostasis were also induced, including the putative thioredoxin reductase *PA14_09950 (dnoP),* despite marked downregulation of the catalase gene *katE*. Notably, genes associated with respiratory metabolism were extensively remodeled, as indicated by the induction of denitrification genes and high-affinity terminal oxidases, along with the repression of low-affinity terminal oxidases. Upregulated denitrification genes included the *nirSMCFLGHJEN* operon that encodes nitrite reductase, the *norCBD* operon encoding the nitric oxide reductase, and the *nosRZDFYL* operon encoding the nitrous oxide reductase. Consistently, terminal oxidase systems were differentially regulated, with induction of the *ccoN2O2Q2P2* operon encoding the high-affinity cytochrome cbb3-2 oxidase, and modulation of the *coxABG* and *cyoABCDE* operons encoding terminal oxidases with lower oxygen affinity. Finally, genes involved in pyruvate fermentation were significantly downregulated, including the Zn-dependent alcohol dehydrogenase (*PA14_36660*).

Together, these transcriptional changes indicate a global reprogramming of *P. aeruginosa* physiology under Zn limitation, characterized by repression of growth- and virulence-associated pathways and activation of stress responses and respiratory adaptation mechanisms.

### Zn starvation promotes the activation of the denitrification pathway

The RNA-seq analysis revealed a marked upregulation of denitrification genes in Zn-starved *P. aeruginosa* (**Fig. 2A**), including genes encoding nitrite (*nir*), nitric oxide (*nor*), and nitrous oxide (*nos*) reductases.

**Fig. 2:**
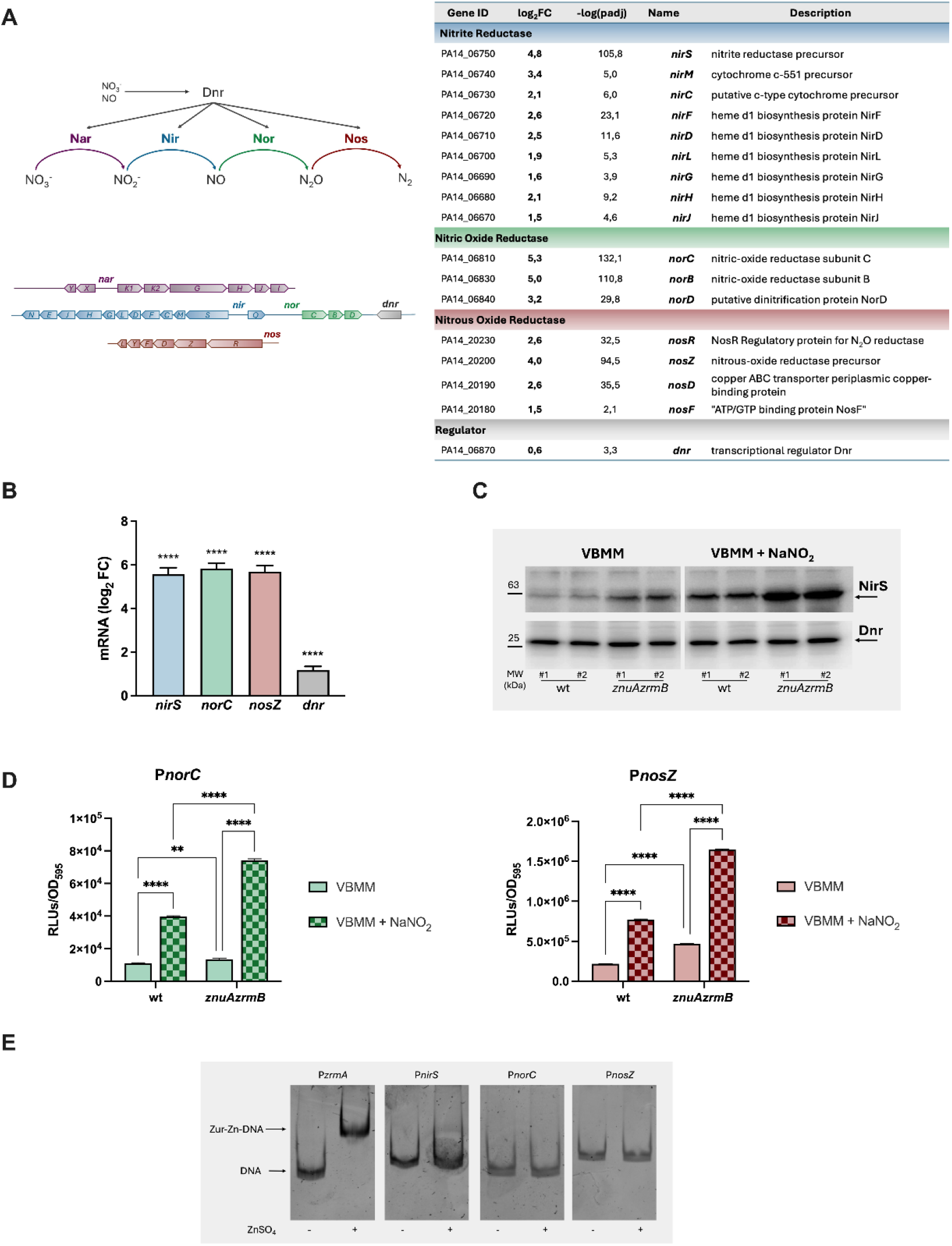
Analyses of the denitrification pathway in Zn-starved *P. aeruginosa*. (**A**) Schematic representation of the denitrification pathway in *P. aeruginosa* and related operons and list of differentially expressed denitrification genes in the *znuAzrmB* strain from the RNAseq analysis. (**B**) RT-qPCR analyses in *znuAzrmB* mutant grown in VBMM. Data are the mean values of three biological replicates ± SD, and statistical analyses were carried out using the Student’s t-test. Asterisks indicate statistical differences between *znuAzrmB* and PA14 wild-type (****p < 0.0001). (**C**) Western blot analysis of NirS and Dnr in PA14 wild-type and *znuAzrmB* mutant strains (two colonies each strain) grown in VBMM supplemented with or without NaNO_2_ 0.1 mM. (**D**) Activity of P*norC* (left panel) and P*nosZ* (right panel) promoters, measured by a lux reporter assay, in PA14 wild-type and *znuAzrmB* mutant strains grown in VBMM supplemented with or without NaNO_2_ 0.1 mM. Bars are the mean values of three biological replicates ± SD, and statistical analyses were carried out using the Two-way ANOVA and Tukey’s multiple comparisons test (**p < 0.01; ****p < 0.0001). **(E)** Electrophoretic mobility shift assays (EMSA) using purified Zur protein and the promoters *nirS, norC* and *nosZ*. Each mixture was treated with 30 µM ZnSO_4_, or untreated, as indicated. Arrows indicate the unbound promoters (DNA) or promoter bound by the Zn cofactored-Zur (Zur-Zn-DNA).

To validate these findings, we compared the expression of key denitrification genes in the two strains in VBMM (**Fig. 2B**). Transcript levels of *nirS*, *norC* and *nosZ*, were significantly higher in the *znuAzrmB* mutant strain than in the wild-type in Zn-restricted conditions. A modest increase in the expression of the regulatory gene *dnr* was also observed.

Since denitrification gene expression is influenced by the availability of alternative electron acceptors, we next examined the effect of nitrite supplementation. Protein expression analysis confirmed that NirS accumulates at higher levels in the Zn-starved mutant than in the wild-type strain, and that its accumulation further increased in the presence of nitrite. In contrast, no difference in Dnr accumulation was observed (**Fig. 2C and Fig S1**). Moreover, we used *lux* transcriptional reporters to monitor *norC* and *nosZ* promoter activity in PA14 wild-type and *znuAzrmB* mutant strains grown in VBMM with or without nitrite. Both promoters displayed higher activity in the *znuAzrmB* mutant compared to the wild-type strain, and their activity was further induced upon nitrite addition (**Fig. 2D**), supporting enhanced activation of the denitrification pathway in the mutant background.

Finally, to assess whether this response is directly controlled by the Zn-responsive regulator Zur, we performed electrophoretic mobility shift assays (EMSA). Whereas Zn-Zur complex bound the control *zrmA* promoter, no binding was detected at the *nirS*, *norC* or *nosZ* promoters (**Fig. 2E**), indicating that Zur does not directly regulate these operons in *P. aeruginosa*.

Together, these data demonstrate that severe Zn limitation promotes the expression of denitrification genes, although this response does not appear to be mediated directly by Zur.

### Zn starvation remodels aerobic respiration in the *znuAzrmB* mutant

The RNA-seq analysis revealed differential regulation of terminal oxidases in the *znuAzrmB* mutant (**Fig. 3A**). Specifically, genes encoding low-affinity terminal oxidases, including the *coxABG* and *cyoABCDE* operons, were downregulated, whereas the *ccoN2O2Q2P2* operon encoding the high-affinity cytochrome cbb3-2 oxidase was upregulated. These transcriptional changes were validated by RT-qPCR analysis in PA14 wild-type and *znuAzrmB* mutant strains grown in VBMM (**Fig. 3B**).

**Fig. 3:**
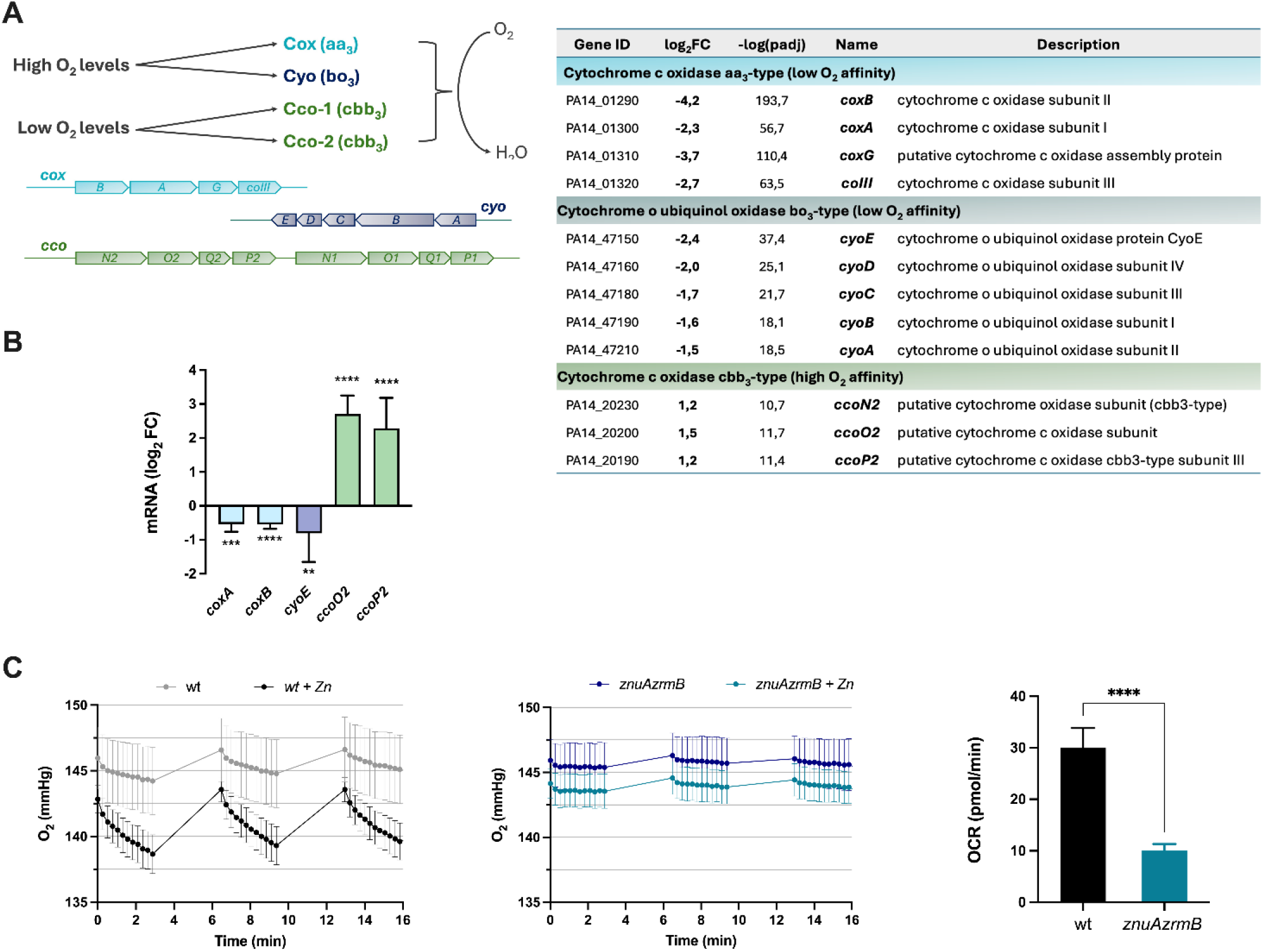
Analyses of aerobic respiration in Zn-starved *P. aeruginosa*. (**A**) Schematic representation of the *P. aeruginosa* terminal oxidases for aerobic respiration and a list of differentially expressed terminal oxidase genes in the *znuAzrmB* strain from the RNAseq analysis. (**B**) RT-qPCR analyses in the *znuAzrmB* mutant grown in VBMM. Data are the mean values of three biological replicates ± SD, and statistical analyses were carried out using the Student’s t-test. Asterisks indicate statistical differences between *znuAzrmB* and PA14 wild-type (**p<0.01; ***p<0.001; ****p < 0.0001). (**C**) Left and middle panels: oxygen levels detected by Seahorse in the growth medium of PA14 wild-type and *znuAzrmB* mutant strains grown in VBMM with or without Zn supplementation, as indicated in the legends. Each group of measures was alternated with 3 minutes of mixing (to promote gas exchange and restore initial oxygen levels). Right panel: oxygen consumption rate (OCR), extrapolated from the third group of measures of the oxygen levels, comparing wt and *znuAzrmB* strains grown without Zn supplementation (Fig S2). Data points represent the average of the eight technical replicates in a representative experiment ± SD. Statistical analysis was performed with the Student’s t-test (****p<0.0001).

To assess whether the transcriptional remodeling of terminal oxidases translated into functional changes in aerobic respiration, oxygen consumption was measured using Seahorse analysis. Under Zn-limiting conditions, only minor differences in oxygen consumption were observed between the *znuAzrmB* mutant and the wild-type strain (**Fig. 3C and S2**). In contrast, Zn supplementation markedly increased oxygen consumption in the wild-type, whereas no such response was observed in the *znuAzrmB* mutant consistent with its inability to efficiently import Zn (**Fig 3C**). As a consequence, a pronounced difference emerged between the two strains under Zn-replete conditions, with a markedly reduced oxygen consumption by the *znuAzrmB* mutant compared to the wild-type strain, indicative of a reduced respiratory activity (**Fig. 3C**). The reduced oxygen utilization observed in the mutant is consistent with the transcriptional reprogramming of terminal oxidases identified by RNA-seq and RT-qPCR analyses. Together, these findings indicate that Zn limitation leads to a remodeling of aerobic respiration in *P. aeruginosa*, characterized by altered terminal oxidase expression and reduced oxygen utilization.

### Zn starvation affects membrane-associated functions and enhances tolerance to aminoglycosides

The shift toward denitrification and the remodeling of aerobic respiration are expected to affect cellular energy production. To determine whether the observed changes in respiratory pathways impact cellular energetics, ATP levels were quantified in PA14 wild-type and *znuAzrmB* mutant cells grown under Zn-limited conditions. The mutant strain exhibited significantly lower ATP levels compared to the wild-type, indicating reduced cellular energy (**Fig. 4A**).

**Fig. 4.**
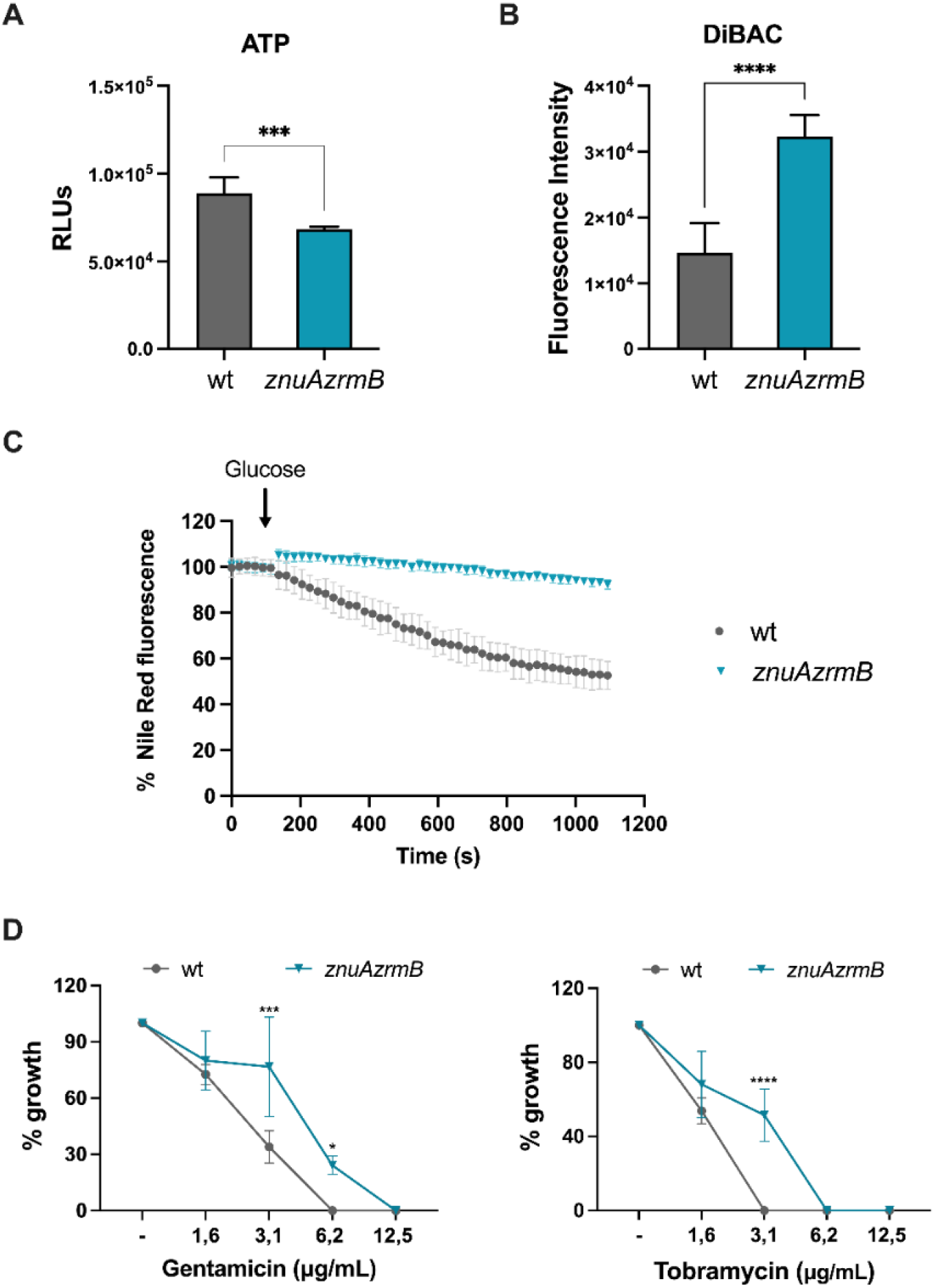
Modification of ATP levels, membrane depolarization, efflux activity and aminoglycoside tolerance in Zn-starved *P. aeruginosa*. (A) Quantification of ATP in PA14 wild-type and *znuAzrmB::FLP* strains, using the BacTiter-Glo kit, where the luminescence (RLUs) is proportional to the number of ATP molecules. **(B)** Membrane depolarization of PA14 wild-type and *znuAzrmB* strains, as assessed by DiBAC_4_(3) fluorescence. In both panels, each bar represents the mean ± SD of two biological replicates deriving from two independent experiments, each as a technical triplicate. Statistical analyses in (A) and (B) were performed by an unpaired Student’s t-test (***p<0.001; ****p < 0.0001). (**C)** Nile Red efflux in PA14 wild-type and *znuAzrmB* strains following supplementation with glucose 3.25 mM (time point 90 s) as indicated by the arrow. Fluorescence was recorded at regular intervals as described in Materials and Methods section. (**D**) Growth of *P. aeruginosa* strains in VBMM + 2,5 μ ϵDTA with increasing concentration of gentamicin (left) or tobramycin (right), normalized to the untreated control (set to 100%). Statistical significance was assessed using two-way ANOVA with Sidak’s multiple-comparisons test. Each point represents the mean ± SD of two biological replicates. Asterisks indicate differences between the strains at the same antibiotic concentration: ****p < 0.0001; ***p < 0.001; *p < 0.05.

Because aerobic respiration is a major contributor to the proton-motive force (PMF), we next evaluated whether Zn starvation could affect membrane potential. Using the potential-sensitive dye DiBAC₄(3), we observed substantially higher fluorescence in the *znuAzrmB* mutant, indicating an enhanced membrane depolarization, consistent with a reduced PMF (**Fig. 4B**).

To determine whether these energetic changes influence membrane-associated transport processes, we assessed efflux activity using a Nile Red efflux assay. The *znuAzrmB* mutant showed reduced efflux activity compared to the wild-type (**Fig. 4C**), which is consistent with both the transcriptional downregulation of efflux-associated genes (**Table S1**) and the reduced ATP levels and membrane potential observed in the mutant **(Fig. 4A-B).** This finding suggests a broader impairment of energy-dependent transport systems under Zn-limited conditions.

Next, we examined the impact of these physiological changes on tolerance to aminoglycosides, whose uptake depends on the PMF generated by respiratory metabolism. A reduction in PMF is expected to limit intracellular drug accumulation. Consistent with this hypothesis, growth inhibition assays showed that the *znuAzrmB* mutant tolerated higher concentrations of aminoglycosides than the wild-type strain (**Fig. 4D**). Specifically, growth of the wild-type was inhibited at 6.2 µg/mL gentamicin and 3.1 µg/mL tobramycin, whereas inhibition of the *znuAzrmB::FLP* mutant required higher concentrations (12.5 µg/mL gentamicin and 6.2 µg/mL tobramycin).

Together, these results indicate that Zn starvation-induced metabolic remodeling reduces cellular energy status and PMF, impairs membrane transport processes, and enhances tolerance to aminoglycosides.

### Zn starvation induces a redox imbalance

Transcriptomic analysis of the *znuAzrmB* mutant revealed induction of genes associated with oxidative stress responses. To evaluate the intracellular redox state under Zn starvation, we compared oxidative stress-related parameters in *P. aeruginosa* wild-type and *znuAzrmB* mutant strains.

Consistent with the transcriptomic data, the mutant strain exhibited significantly increased intracellular reactive oxygen species (ROS) levels together with a reduced glutathione (GSH) pool, indicative of redox imbalance under Zn-limited conditions (**Fig. 5A** and **5B**).

**Fig. 5.**
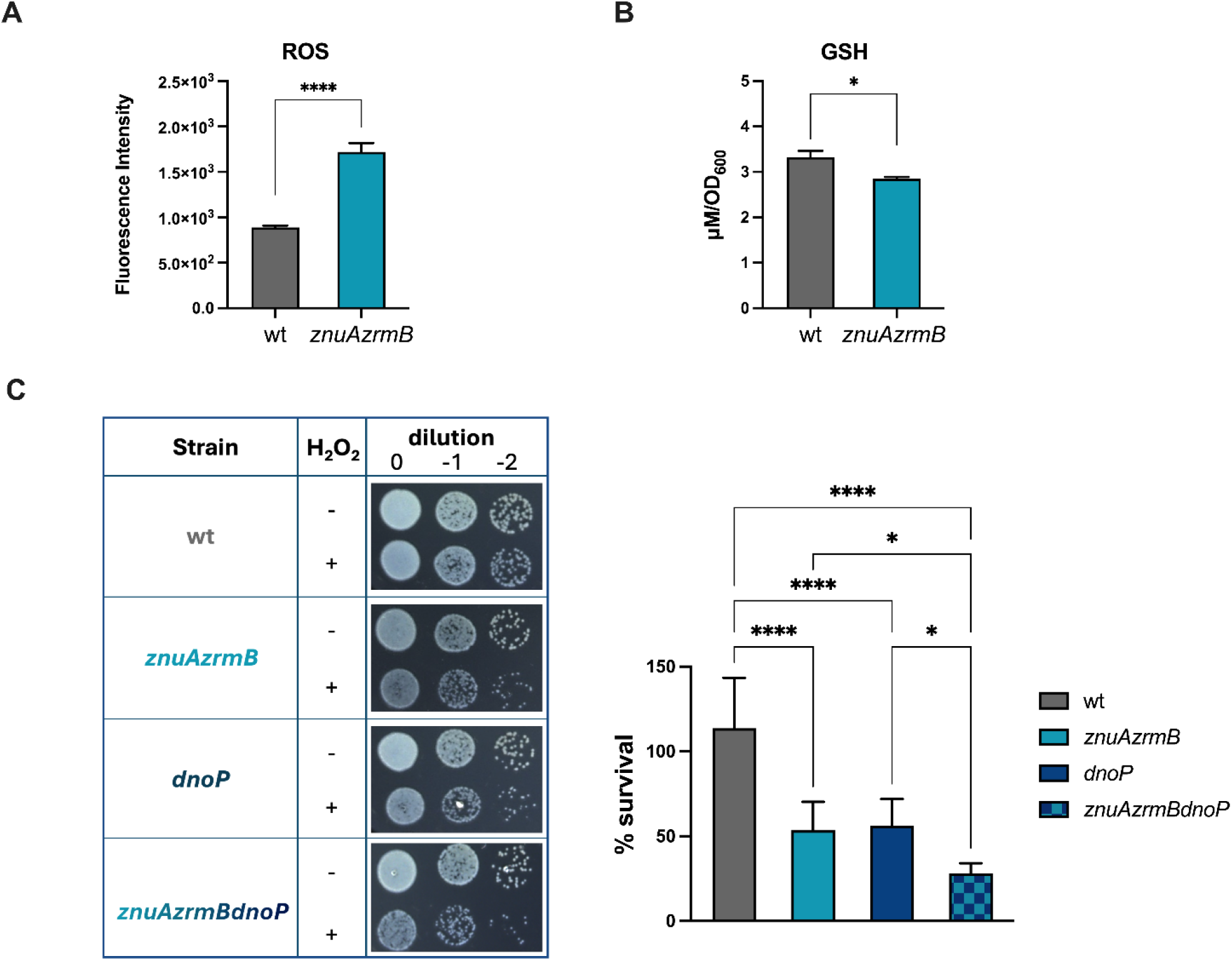
Oxidative stress responses in Zn-starved *P. aeruginosa*. (A) Measurement of intracellular reactive oxygen species levels in PA14 wild-type and *znuAzrmB* mutant strains detected by H_2_DCFDA fluorescence (Ex 485 nm/Em 535 nm). Data represent the mean ± SD of two independent biological replicates analyzed in technical triplicate. Statistical significance was determined using unpaired t-test (****p< 0.0001). (B) Intracellular reduced glutathione levels in PA14 wild-type and *znuAzrmB* mutant strains were quantified using DTNB (Ellman’s reagent) by measuring absorbance at 415 nm. Data represent the mean ± SD of two independent biological replicates analyzed in technical triplicate. Statistical significance was determined using unpaired t-test (*p < 0.05). (C) Hydrogen peroxide sensitivity of PA14 wild-type and mutant strains. Images show serial dilutions of PA14 wild-type, *znuAzrmB*, *dnoP* and *znuAzrmBdnoP* mutant strains spotted on LB plates following exposure to 0.75 mM H₂O₂ (left panel). Survival percentages were calculated using the CFU counts from untreated samples as 100% (right panel). Values represent the mean of four biological replicates ± SD. Statistical analysis was performed using one-way ANOVA with Tukey’s multiple comparison test (*p < 0.05; ****p < 0.0001).

Among the most strongly induced genes identified by RNA-seq was *dnoP*, which encodes a recently characterized thioredoxin reductase (**Fig. 1** and **Table S1**) (21). Deletion of *dnoP* significantly reduces resistance to hydrogen peroxide, with an additive effect in the *znuAzrmBdnoP* mutant (**Fig. 5C**), suggesting a critical role for this thioredoxin-based system in protecting *P. aeruginosa* against oxidative stress during Zn limitation.

### Zn starvation is associated with alterations in quorum sensing signaling

Transcriptomic analysis revealed a reduction, although not reaching statistical significance, in the expression of the QS regulatory genes *lasR* and *rhlR*, accompanied by a strong downregulation of multiple QS-controlled virulence genes, including *lasA*, *lasB*, *aprA*, and *phz* operons in the *znuAzrmB* mutant compared to PA14 (**Table S1**).

To assess the impact of Zn limitation on *P. aeruginosa* QS systems, we measured the levels of major QS signaling molecules produced by the PA14 wild-type and the *znuAzrmB* mutant. Levels of N-butyryl-L-homoserine lactone (C_4_-HSL) and alkylquinolones (AQs) were significantly reduced in the *znuAzrmB* mutant, whereas N-3-oxododecanoyl homoserine lactone (3OC_12_-HSL) levels remained unchanged (**Fig. 6**). These findings indicate that Zn starvation selectively alters QS output at the metabolite level, consistent with the observed repression of QS-dependent virulence genes.

**Fig. 6:**
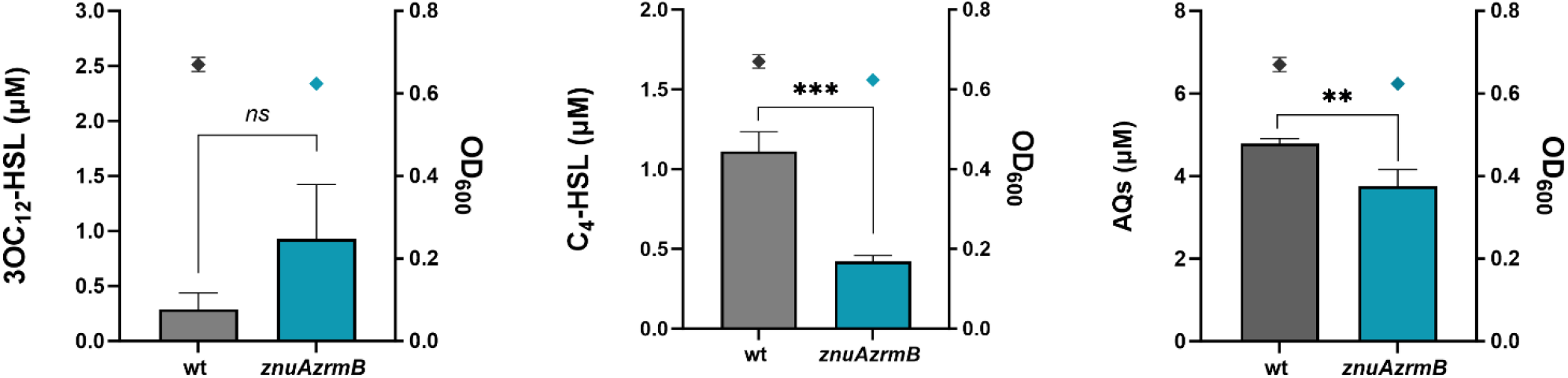
Production of quorum sensing signal molecules in *P. aeruginosa* supernatants. 3OC_12_-HSL (**A**), C_4_-HSL (**B**) ed AQs (**C**) signal molecules quantified by biosensor-based assays in supernatants of PA14 wild-type and the *znuAzrmB* mutant strain. The cell density (OD_600_) values of the PA14 wild-type and *znuAzrmB* cultures at the time point when the supernatants were collected are shown as black and blue diamonds, respectively. Data represent the mean ± SD of three biological replicates. Statistical analysis was performed by Student’s t-test and asterisks indicate statistically significant differences between the strains (**p < 0.01; ***p < 0.001). *ns:* not significant.

## Discussion

Zn limitation represents a major host-imposed stress that bacteria must overcome to persist during infection. In *P. aeruginosa*, adaptation to Zn starvation has been extensively characterized in terms of the Zur regulon. However, the global physiological response to severe Zn deficiency remains incompletely defined. By comparing the transcriptomic profiles of the PA14 wild-type strain and a *znuAzrmB* mutant lacking the major Zn import systems, we provide a comprehensive view of this adaptive response. Our results show that Zn limitation induces a widespread reprogramming of bacterial physiology, characterized by reduced central metabolism, respiratory remodeling, altered redox homeostasis, and attenuation of virulence-associated functions. Together, these changes promote a shift toward a low-energy, stress-adapted state that recapitulates key features of persistence-associated phenotypes.

A notable finding of this study is the substantial overlap between the transcriptional profile of the Zn-starved *znuAzrmB* mutant and those reported for *P. aeruginosa* populations isolated from CF airways (2,4,16,22). Chronic infection in the CF lung is characterized by extensive metabolic rewiring that supports persistence under nutrient limitation, oxidative stress, and restricted oxygen availability. Notably, several of these hallmarks are recapitulated under Zn starvation. In particular, the repression of carbohydrate uptake and central carbon metabolism observed in the mutant is consistent with the reduced reliance on glucose reported in clinical isolates (23) and reflects a reduction in energy-generating and biosynthetic processes required for growth. Moreover, the observed downregulation of the genes enriched in chemotaxis, motility, and attachment, as well as flagellar components, aligns with a non-motile phenotype frequently observed in chronic *P. aeruginosa* infections, where motility components are often downregulated (16). This profile is consistent with previous studies demonstrating that impaired Zn import negatively affects flagellar-dependent bacterial motility (9,24). However, unlike CF isolates, which frequently exhibit enhanced biofilm formation to survive within the thick mucus of the CF lung (16,25), the Zn-starved mutant shows reduced expression of biofilm-associated genes, consistent with previous observations (9).

Additionally, the Zn-starved mutant shows induction of organic sulfur acquisition pathways, which parallels transcriptional adaptations observed in the CF mucus environment, where organosulfonates represent a major sulfur source due to the presence of mucins (26,27). Sulfur metabolism is critical for bacterial survival, supporting the synthesis of amino acids, enzyme cofactors, and antioxidants such as glutathione and thioredoxin, particularly under oxidative stress (28,29). Consistent with this, the *znuAzrmB* mutant showed broad induction of oxidative stress–response genes, suggesting that enhanced sulfur assimilation may help maintain redox homeostasis under Zn limitation. Accordingly, sulfur starvation itself has been shown to trigger oxidative stress responses in *P. aeruginosa* (27,30).

Zn homeostasis is functionally linked to bacterial redox control, as Zn can form coordination complexes with cysteine thiol groups, thereby stabilizing proteins and protecting them from oxidative damage (31). Moreover, impairment of Zn acquisition markedly increases susceptibility to H₂O₂-mediated toxicity (24,32,33). Consistently, we showed that the Zn-starved mutant accumulates reactive oxygen species and exhibits reduced glutathione, indicating a redox imbalance. Additionally, the *znuAzrmB* mutant shows a pronounced induction of *dnoP*, encoding a recently characterized thioredoxin-associated reductase involved in protection against redox-active toxic compounds, such as the fungal gliotoxin produced by *A. fumigatus* during co-infections (21). Interestingly, gliotoxin itself elicits a Zn-starvation response in *P. aeruginosa*, likely through metal chelation, further linking oxidative stress responses to perturbation of Zn homeostasis. Consistently, *dnoP* was also upregulated in PA14 wild-type grown under Zn-limited conditions compared to Zn-supplemented conditions (14), further supporting its role as part of the Zn-starvation response. The increased sensitivity of the *dnoP* mutant to hydrogen peroxide, particularly in the mutant with reduced Zn-uptake ability, supports a role for this system in maintaining redox balance under metal limitation. Despite the activation of oxidative stress responses, the catalase gene *katE* is markedly downregulated. Unlike KatA and KatB, the role of KatE remains unclear and does not appear to involve H₂O₂ detoxification (34).

One of the most prominent features of the Zn-starved transcriptome is the extensive remodeling of respiratory metabolism. Genes involved in denitrification and high-affinity terminal oxidases were strongly induced, whereas low-affinity terminal oxidases were repressed, consistent with a shift toward microaerobic and anaerobic respiratory pathways despite aerobic growth conditions. Importantly, our experimental validation demonstrates that these transcriptional changes are accompanied by functional alterations in respiratory activity upon Zn supplementation. Oxygen consumption was reduced in the Zn-starved mutant, and the difference became particularly evident upon Zn supplementation, which increased oxygen utilization in the wild-type but not in the Zn-uptake-deficient strain. This finding supports a role for intracellular Zn availability as a previously unrecognized factor influencing respiratory activity in *P. aeruginosa* and suggest that impaired Zn acquisition triggers a respiratory adaptation characterized by reduced oxygen utilization and remodeling of terminal oxidase expression. Consistent with this model, the mutant exhibited increased expression of denitrification enzymes and enhanced nitrite-dependent induction of denitrification pathways. A similar response has been observed in *P. denitrificans*, where Zn starvation strongly induces denitrification genes (35). However, unlike in *P. denitrificans*, this response does not appear to be directly controlled by the Zn-responsive regulator Zur, indicating that respiratory remodeling under Zn limitation is mediated through indirect regulatory mechanisms in *P. aeruginosa*. The molecular basis of this response remains unclear. It may involve alterations in cellular redox homeostasis, changes in global regulatory networks, or secondary effects arising from the reduced activity of Zn-dependent metabolic functions. Further studies will be required to distinguish among these possibilities and define how Zn availability is coupled to respiratory regulation.

Interestingly, concurrent induction of Zn-starvation responses and anaerobic metabolism has often been reported, including in *P. aeruginosa* co-cultured with *A. fumigatus*, where denitrification genes are induced alongside Zn-starvation responses (21), as well as under host-relevant conditions, including *P. aeruginosa* CF sputum-derived transcriptomes, which display concurrent signatures of Zn restriction and oxygen limitation (4). While Zn restriction is a well-established component of nutritional immunity and anaerobic respiration is typically attributed to hypoxic niches within infected tissues, the induction of an anaerobic respiratory program in the *znuAzrmB* mutant under aerobic growth conditions suggests that Zn limitation itself can contribute to respiratory reprogramming independently of environmental oxygen availability.

Consistently, multiple genes associated with anaerobic metabolism are induced, including the anaerobically induced porin gene *oprE*, *nnrS* (*PA14_29660/PA2662*), encoding a membrane protein required for efficient growth under denitrification conditions (36), and *ccpR*, encoding a cytochrome c peroxidase within the ANR regulon (37). In addition, *nrdG* is induced, consistent with activation of class III ribonucleotide reductase systems operating under oxygen-limited conditions, such as those encountered in deeper biofilm layers (38).

This respiratory adaptation appears to have an energetic cost. The Zn-starved mutant exhibits reduced energy production, likely reflecting both the lower bioenergetic efficiency of denitrification compared with aerobic respiration (39) and the concomitant downregulation of TCA cycle genes (**Fig. 1** and **Table S1**). Consistent with this energetic limitation, several ATP-consuming processes are repressed, including flagellar assembly and multidrug efflux systems. Despite this global reduction in energy-demanding processes, ATP production remains essential for viability, as multiple energy-dependent metal uptake systems are strongly induced under Zn limitation. Accordingly, increased expression of ATP synthase genes (**Table S1**) may represent a compensatory response to maintain cellular energy homeostasis under conditions of reduced respiratory efficiency. Taken together, these observations support the existence of a metabolic downshift in Zn-starved cells, in which energy-intensive functions are selectively decreased, while processes required for survival and metal acquisition are maintained.

Additionally, the Zn-starved mutant displays membrane depolarization, consistent with a reduced PMF. Given that aminoglycoside uptake depends on PMF (40), this depolarization likely contributes to the increased tolerance to tobramycin and gentamicin, in agreement with multiple studies linking central carbon metabolism and respiratory activity to aminoglycoside susceptibility (41–44). For instance, fumarate-driven stimulation of TCA cycle activity enhances respiration, increases PMF, and promotes aminoglycoside uptake and killing, whereas glyoxylate metabolism attenuates respiratory flux, reduces PMF, and decreases antibiotic efficacy (45). Notably, increased reliance on the glyoxylate shunt is a hallmark of chronic *P. aeruginosa* adaptation in the CF lung (4) and may contribute to intrinsic aminoglycoside tolerance. Moreover, clinical isolates from CF patients most commonly exhibit reduced intracellular accumulation, often referred to as membrane impermeability (46). This phenotype is typically associated with an increased efflux (e.g., via MexXY-OprM) (47) and decreased drug influx under conditions of low respiratory activity or anaerobiosis (48,49). In the Zn-starved mutant, efflux-associated genes are overall reduced, consistent with decreased activity as confirmed by Nile Red assays. These observations suggest a mechanism primarily driven by reduced intracellular uptake rather than increased efflux or enzymatic inactivation, although further investigation is required to conclusively establish the resistance mechanism.

QS signaling is selectively altered under Zn starvation. Although transcriptional changes in QS regulators are modest, production of key signaling molecules, including C_4_-HSL and AQs, is significantly reduced, accompanied by repression of multiple QS-dependent virulence factors, including Zn-dependent extracellular proteases and phenazine biosynthesis, in agreement with previous evidence (24,50–52).

In chronic *P. aeruginosa* infections, *lasR* loss-of-function mutations are frequently selected (16,22,53,54). These variants exhibit reduced oxygen consumption, altered respiratory metabolism, and increased tolerance to aminoglycosides and oxidative stress (54). Consistent with these phenotypes, QS plays a central role in coordinating respiratory pathways by regulating terminal oxidases and denitrification genes: C_4_-HSL and 3OC_12_-HSL repress denitrification, whereas PQS modulates this process both transcriptionally and post-transcriptionally through iron chelation (55–57). In parallel, QS promotes expression of aerobic terminal oxidases, including *coxAB* and *cyoABCD* (58).

In this context, the Zn-starved mutant displays a QS signaling profile characterized by reduced C_4_-HSL and AQs production, whereas 3OC_12_-HSL levels remain unchanged. Consistently, Zn supplementation has been reported to stimulate *lasR* expression and the RhlR-dependent genes *rhlA* and *rhlB*, whereas Zn limitation impairs PQS and rhamnolipid production (15,50). Although our data do not establish a causal relationship between altered QS signaling and respiratory remodeling, given the central role of QS in balancing aerobic and anaerobic respiratory programs, these observations raise the possibility that Zn limitation may indirectly influence respiratory adaptation by modulating QS signaling. Alternatively, respiratory rewiring under Zn limitation may arise independently of QS alterations and instead represent a compensatory response to limit ROS generation associated with aerobic respiration. Clarifying the interplay between Zn homeostasis, QS signaling, and respiratory regulation will require further investigation and lies beyond the scope of the present study.

Taken together, our findings support a model in which Zn limitation drives a coordinated physiological transition characterized by reduced metabolic activity, respiratory reprogramming, redox imbalance, and altered regulatory signaling. This state recapitulates several hallmarks of the adaptive phenotype of *P. aeruginosa* in chronic CF infections, suggesting that metal limitation may represent an important environmental cue contributing to the emergence of chronic infection-associated phenotypes. The extensive physiological changes triggered by Zn limitation also provide a mechanistic framework to interpret recent transcriptome-based studies aimed at improving laboratory models of cystic fibrosis infection. By comparing *P. aeruginosa* transcriptomes obtained *in vitro* with those recovered directly from CF sputum, Lewin *et al.* showed that incorporation of Zn limitation markedly increased the similarity between laboratory-grown bacteria and the *in vivo* transcriptome, identifying Zn availability as a critical environmental parameter shaping bacterial physiology in the CF lung (59). Our results extend these observations by showing that Zn limitation not only alters the expression of Zn homeostasis genes but also reshapes respiratory metabolism, redox homeostasis, quorum sensing, and antibiotic tolerance, providing a mechanistic explanation for why Zn limitation improves the physiological and transcriptional resemblance of laboratory-grown *P. aeruginosa* to bacteria growing in the CF lung.

From a clinical perspective, these results highlight a potential consequence of host-imposed nutritional immunity. By restricting Zn availability, the host may promote bacterial states that are less metabolically active yet more tolerant to antibiotic treatment. This raises important considerations for therapeutic strategies targeting metal homeostasis, as well as for the use of aminoglycosides in environments where metal limitation and low respiratory activity prevail.

Overall, this work provides a comprehensive framework linking Zn homeostasis to global metabolic regulation, respiratory adaptation, and antibiotic tolerance in *P. aeruginosa*. Further studies dissecting the regulatory circuits connecting Zn sensing, redox balance, and quorum sensing will be essential to fully understand how metal availability shapes bacterial physiology during infection.

## Materials and Methods

### Reagents

All chemicals were purchased from Sigma Aldrich unless otherwise specified.

### Bacterial Strains and Growth Conditions

Bacterial strains used in this work are listed in Table S2 and were plated on *Pseudomonas* Isolation Agar (PIA) or Luria-Bertani agar (tryptone 10 g L^−1^, yeast extract 5 g L^−1^, NaCl 10 g L^−1^) supplemented with antibiotics when needed (for *E. coli*, gentamicin 10 μg ml^-1^, ampicillin 100 μg ml^-1^, or kanamycin 50 μg ml^-1^; for *P. aeruginosa*, gentamicin 100 μg ml^-1^) and incubated at 37 °C. *E. coli* and *P. aeruginosa* liquid cultures were routinely grown in LB at 37 °C under shaking. For Zn-starvation conditions, Vogel-Bonner Minimal Medium (VBMM: MgSO_4_-7H_2_O 0,192 g L^-1^, citric acid 2 g L^-1^, anhydrous K_2_HPO_4_ 10 g L^-1^, NaNH_4_HPO_4_ - 4H_2_O 3.5 g L^-1^, glucose 2 g L^-1^, FeSO_4_ 2 μM) supplemented or not with EDTA 5 µM (E-VBMM). To minimize Zn contamination, VBMM was prepared in disposable plastic containers and sterilized by filtration through Vacuum Filter-Storage Bottle Systems (0.22 μm, Corning), as previously described (51).

### RNA Extraction, Reverse Transcription and Real-Time qPCR

Overnight LB cultures of three independent colonies of *P. aeruginosa* PA14 were diluted 1:1000 in VBMM or E-VBMM, and total RNA was extracted from exponentially growing cultures after treatment with RNAprotect (Qiagen). RNA extraction was performed using the RNAeasy kit (Qiagen) according to the manufacturer’s protocol, with the addition of DNase (Qiagen) and using lysozyme.

RNA concentration was determined with a NanoDrop™ Lite Spectrophotometer (Thermo Fisher Scientific). From each sample, 1 µg of RNA was reverse transcribed with the PrimeScript RT Reagent Kit and gDNA Eraser (Takara Bio Inc.). The primers used for RT-qPCR were designed using Primer3 (Table S3) (60). RT-qPCR reactions were performed in triplicate in 10 µL reaction mixtures containing cDNA 50 ng, primers 0.3 µM, and 50% PowerUp SYBR Green Master Mix (Thermo Fisher Scientific). Amplifications were performed in a QuantStudio 3 real-time PCR system (Thermo Fisher Scientific) thermocycler with the following parameters: (i) initial denaturation at 95 °C for 4 min; (ii) 40 cycles of denaturation at 95 °C for 20 s, primer annealing at 60 °C for 30 s and extension at 72 °C for 30 s; (iii) melting curve, from 50 to 90 °C (rate: 0.58 °C every 5 s). The mRNA fold induction was calculated using the ΔΔCt method (61) and normalized to *rpsL* housekeeping gene.

### RNAseq sample preparation, reads mapping and gene expression analysis

PA14 wild-type and *znuAzrmB* mutant strain (three colonies each) were grown overnight in LB, diluted 1:1000 in E-VBMM at 37 °C until the exponential phase; cultures were then treated with RNAprotect and RNA extraction was performed. The concentration and purity of RNA were determined using a NanoDrop™ Lite Spectrophotometer (Thermo Fisher Scientific). rRNA depletion and RNA sequencing were performed at IGA Technology Services Srl Service Provider (Udine, Italy). For cDNA libraries, 40 million paired-end reads of 150 bps were generated for each sample. For the RNA-seq experiments, raw reads were first processed using Trimmomatic (v0.36) (62) to remove low-quality bases and adapters. Reads shorter than 35 nucleotides (nt) were discarded. The processed reads were then mapped to the *P. aeruginosa* UCBPP-PA14 genome (NCBI: NC_008463.1) (63) using the Bowtie2 aligner (64) with default parameters. For gene expression analysis, reads mapping to each annotated coding sequence (CDS) in the *P. aeruginosa* UCBPP-PA14 genome were counted using HTSeq-count (v1.99.2) (65). Counts were normalized using the regularized log transformation function (rlogTransformation) in the DESeq2 R package (66), with the blind option set to TRUE. Differential gene expression (DEG) analysis between mutant and wild-type transcriptomes was performed using DESeq2. Genes were considered statistically significant if they exhibited a Log2(Fold Change) ≥ |1.3| and an adjusted p-value ≤ 0.05. DEGs were inspected, and functional class enrichment was performed using an adapted version of the term_enrich.py Python script (available at https://zenodo.org/records/1162703) (4). This script utilized gene-term associations from the COG and KEGG classification databases, obtained from *Pseudomonas*.com. Functional classes with an adjusted p-value (using Bonferroni correction for multiple testing) ≤ 0.05 were considered significantly enriched.

### Construction of promoter-*lux* fusions

Fragments spanning approximately 200-300 base pairs upstream of the coding sequences of *norC* and *nosZ* were amplified from *P. aeruginosa* PA14 genomic DNA (extracted with Quick-DNA Fungal/Bacterial Kit, Zymo Research) by PCR, with Expand^TM^ High-Fidelity DNA Polymerase (Roche Life Science) and primers listed in Table S3. Each fragment was digested with EcoRI and SacI restriction enzymes (New England Biolabs), purified from agarose gel using the Zymoclean Gel DNA Recovery Kit (Zymo Research), and ligated into the pETS*lux* plasmid, as previously described (7,67). *E. coli* DH5α competent cells were transformed with the ligation mixtures, and the plasmids were then isolated from gentamicin-resistant clones using the Plasmid Isolation Kit (Zymo Research). The insertion of promoters was verified by EcoRI/SacI restriction. Plasmids with the promoters-*lux* fusion (Table S2) were transferred to the *P. aeruginosa* PA14 wild-type and *znuAzrmB::*FLP mutant strains by triparental mating, using HB101 pRK2013 as the helper strain. Exconjugants were selected on PIA supplemented with gentamicin 100 mg L^-1^.

### *P. aeruginosa* growth and luminescence analyses

Overnight LB cultures of *P. aeruginos*a strains carrying P*norC*-lux or P*nosZ*-lux plasmids were diluted 1:1000 in E-VBMM supplemented or not with NaNO_2_ 100 µM. Bacterial strains were grown for 18 h at 37 °C, 120 rpm, and the end-point growth and luminescence were measured in a black microplate with a transparent bottom (Greiner Bio-One) by a Sunrise™ microplate reader, simultaneously recording the optical density at 595 nm (OD_595_) and the relative luminescence units (RLUs, integration time 1000 ms). Each experiment was tested in biological and technical triplicate and was repeated at least twice.

### Western Blot

Overnight LB cultures of *P. aeruginos*a strains were diluted 1:1000 in E-VBMM supplemented or not with NaNO_2_ 100 µM and grown 18 h at 37 °C, 120 rpm. Aliquots of 5 × 10^8^ cells were harvested, lysed by resuspending bacteria in sample buffer containing sodium dodecyl sulfate (SDS) and β-mercaptoethanol, and boiled for 5 min at 100 °C. Proteins were run on an SDS-polyacrylamide gel and blotted onto a nitrocellulose membrane (Hybond ECL; Amersham). Ponceau staining was performed to verify equal loading of the gels. Western blot analyses were carried out with rabbit polyclonal antibodies against *P. aeruginosa* NirS and Dnr (68,69) and using a Horseradish Peroxidase (HRP)-conjugated anti-rabbit secondary antibody. Signals were detected by enhanced chemiluminescence (ECL; GE Healthcare) using a Chemidoc imaging system (Bio-Rad Laboratories), and densitometric analysis was carried out with ImageJ.

### Electromobility Shift Binding Assay (EMSA)

The expression and purification of *P. aeruginosa* Zur was performed following a previously described protocol with some modifications, as recently described (24,70). For the EMSA experiments, promoter *nosZ*, *norC*, and *nirS* were obtained by PCR amplification from genomic PA14 DNA using the primers listed in Table S3. The binding assays were performed as recently described, using the *zrmA* promoter as a control (24). Briefly, reactions were performed with a mixture composed of the 5X Zn-less Binding Buffer (Tris 50 mM, KCl 200 mM, MgCl_2_ 50 mM, DTT 5 mM, and glycerol 25%, pH 8.0), 30 ng of DNA, TPEN 30 μM, Zur protein 1 µM in presence or absence of ZnSO_4_ 30 μM and incubated at room temperature for 30 min. Samples were loaded on a 7.5% polyacrylamide native gel containing glycerol 2.5% in Tris Borate Buffer and run at 4 °C. The bands corresponding to free or Zur-bound promoters were visualized by staining the gel with ethidium bromide 0.1% and revealed with UV light using a Chemidoc imaging system (Bio-Rad Laboratories).

### Seahorse analysis

Cellular oxygen consumption rate (OCR) and oxygen levels were measured by the extracellular flux analyzer XFe96 (Seahorse Bioscience) in the Hyp-ACB facility at Sapienza University, as previously described (71). Single colonies of PA14 wild-type and *znuAzrmB* strains grown overnight in LB were inoculated in VBMM supplemented and grown 18h at 37°C, 120 rpm. Bacteria were water-diluted to reach OD_600_ = 0.02, and aliquots of 90 µl were placed in each well of the 96-wells Seahorse cell culture plate precoated with collagen (72). Plates were centrifuged for 10 min at 4000 rpm to allow bacteria to adhere to the surface. After centrifugation, 90 µl of 2X VBMM was added in each well. The concentration of cells used in the experiment was further verified by crystal violet assay, by measuring the OD_600_ after solubilization of the dye; the OD_600_ of the stained seeded bacteria was found to be linearly dependent on the OD_600_ of the corresponding liquid growth. Each sample was analyzed at 37°C in up to eight wells per experiment, and two independent experiments were carried out; VBMM controls were included in at least two wells per experiment.

### ATP measurement

ATP was measured from metabolically active *P. aeruginosa* cells grown in VBMM using BacTiter-Glo™ Microbial Cell Viability Assay (Promega). Briefly, PA14 wild-type and *znuAzrmB* strains were grown to a logarithmic phase in VBMM, and the cell cultures were normalized to OD_595_ 0.5. The BacTiter-Glo™ Reagent was reconstituted according to manufacturer instructions and equilibrated at room temperature for 15 min. Equal volumes of BacTiter-Glo™ Reagent and cell culture were added to a 96-well black microplate (Greiner, Bio-One). The mixtures were agitated briefly on an orbital shaker and incubated for 5 min in the dark. The RLUs (integration time 1000 ms) were recorded by a Sunrise™ microplate reader (Tecan). Background luminescence, determined from wells containing only VBMM mixed 1:1 with BacTiter-Glo™ Reagent, was subtracted from the luminescence readings of each well.

### Membrane depolarization assay

The assay for measuring cytoplasmic *P. aeruginosa* membrane depolarization was based on a previously described assay (73). Cultures of *P. aeruginosa* were grown in E-VBMM up to a logarithmic phase, at which time cells were exposed to the membrane potential-sensitive dye bis-(1,3-dibutylbarbituric acid) trimethine oxonol [DIBAC_4_(3); Thermo Fischer Scientific] 10 µg mL^-1^ at 37 °C for 5 min in the dark. Bacteria were then pelleted by centrifugation (3000 rpm) and resuspended in PBS. After 20 min at 37 °C, samples were dispensed in a 96 well black microplate in technical triplicates and membrane depolarization-dependent fluorescence emitted by cells was measured by a Sunrise™ microplate reader (λ_ex = 490 nm, λ_em = 518 nm). Each experiment was performed on two independent colonies and was repeated three times.

### Nile Red Efflux Assay

Nile Red efflux was measured following a previously established protocol, with minor modifications (74). Briefly, overnight LB cultures of PA14 wild-type and the *znuAzrmB* mutant were diluted 1:1000 in E-VBMM and grown overnight at 37 °C with shaking. Cells were harvested, washed twice, and resuspended in potassium phosphate buffer (PPB; KPO₄ 20 mM, MgCl₂ 1 mM) to an OD₆₀₀ of 1, followed by incubation at room temperature for 15 min. Cells were de-energized by adding carbonyl cyanide m-chlorophenylhydrazone (CCCP) 10 µM and incubating at 35 °C for 20 min with shaking (140 rpm). Cells were treated with Nile Red 5 µM and incubated at 37 °C for 1.5 h (140 rpm) to allow dye accumulation, then equilibrated at room temperature for 30 min. Cells were pelleted, washed, resuspended in PPB, and aliquoted into a black 96-well microplate (Greiner Bio-One) in technical triplicate. Fluorescence (λ_ex = 544 nm, λ_em = 650 nm) was recorded at 20 s intervals. After baseline measurement, 3.25 mM glucose was added, and fluorescence was registered over time. Data were expressed as percentage fluorescence relative to untreated controls over time. The experiment was performed in technical and biological triplicate and was repeated twice.

### *P. aeruginosa* growth assay under antibiotic treatment

*P. aeruginosa* wild-type and *znuAzrmB::FLP* mutant strain (Table S2) grown overnight in LB medium were diluted 1:2000 in VBMM with EDTA 2,5 μM, with or without increasing concentrations of gentamycin or tobramycin (1.6, 3.1, 6.2, 12.5 μg mL^-1^). A volume of 0.2 mL of each sample was dispensed in a 96-microwell (Greiner Bio-One), incubated at 37 °C in a Sunrise™ microplate reader (Tecan) with periodic shaking. OD_595_ after 18 h was registered and normalized to the untreated control (set to 100%). Each sample was tested in three biological and technical replicates.

### Measurement of Intracellular ROS levels

Intracellular ROS was measured following an already described protocol (75) with minor modifications. Overnight LB inocula of two independent colonies of PA14 wild-type and *znuAzrmB* mutant were inoculated 1:1000 in VBMM and grown 37 °C with shacking until reaching an OD_600_ = 0.5. Cells were then washed twice with PBS and incubated with H_2_DCFDA 10 μM or DMSO (control) for 20 min at 37 °C in the dark. Subsequently, 0.2 mL of each sample was dispensed in a black 96-microwell (Greiner Bio-One) in technical triplicate and fluorescence (λ_ex = 485 nm, λ_em = 535 nm) was recorded using a Sunrise™ microplate reader (Tecan). The experiment was repeated three times.

### Quantification of intracellular GSH

Intracellular reduced glutathione levels were measured using Ellman’s reagent [5,5′-dithiobis-(2-nitrobenzoic acid), DTNB], following a previously described protocol (76) with some modifications. Specifically, overnight LB cultures of two independent colonies of PA14 wild-type and the *znuAzrmB* mutant were diluted 1:1000 in VBMM and grown at 37 °C with shaking until reaching the exponential growth phase. Cells corresponding to 1 OD unit were collected, centrifuged 5,000 × *g* for 10 min at 4 °C, washed with cold PBS, and centrifuged again. Pellets were resuspended in PBS containing EDTA 5 mM and metaphosphoric acid (MPA) 5% and incubated on ice for 20 min. Cells were disrupted by repeated cycles of freeze/thawing and after centrifuging at 10,000 × *g* for 15 min at 4 °C, supernatants were neutralized by the slow addition of KOH 0.3 mM.

For glutathione quantification, 50 μL of each sample was mixed with 150 μL of phosphate buffer (100 mM, pH 7.5, EDTA1 mM) containing DTNB 0.1 mM in a 96-well microplate (Sarstedt). Absorbance at 415 nm was measured after incubation for 15 min at 37 °C. A standard curve was generated using reduced glutathione concentrations from 1 to 50 μM. Each sample was analyzed in technical triplicate, and the experiment was repeated twice.

### *P. aeruginosa dnoP* mutant construction

The *dnoP* deletion mutant was obtained using the gene replacement method with minor modifications (51,77). The gentamicin resistance cassette was obtained by BamHI (New England Biolabs) digestion of plasmid pSP856 (Table S2). The 5’ and the 3’ terminal fragments of *dnoP* were amplified using PA14 wild-type DNA as a template and with the primers listed in Table S3. The EcoRI/BamHI and BamHI/HindIII (New England Biolabs) digested 5’ and 3’ fragments, and the gentamicin resistance cassette was cloned into plasmid pEX18Tc. The resulting plasmid was mobilized to PA14 wild-type by tri-parental mating and the *dnoP* deletion in PA14 was confirmed by PCR using the primers in Table S3.

### Hydrogen peroxide tolerance assay

Hydrogen peroxide tolerance assay was performed as previously described (24). Briefly, PA14 wild-type and mutant strains were inoculated 1:1000 in E-VBMM and incubated at 37 °C overnight with shaking. Bacteria were diluted to 10^6^ CFU mL^-1^ in sterile PBS and exposed to H_2_O_2_ 0.75 mmol L^-1^ or left untreated. The samples were incubated at 37 °C. After 1 h, 1000 units of catalase were added to inactivate the hydrogen peroxide, and serial dilutions of bacteria were plated on LB-agar for CFU counting. The number of colonies in each spot was quantified using an automatic colony counter (Scan500 ®, Interscience). The survival percentage was calculated by the formula: (CFU of exposed bacteria / CFU of unexposed bacteria) x 100.

### Quorum sensing molecules measurement

For the assessment of quorum sensing (QS) molecule production, overnight LB cultures of *P. aeruginosa* strains were diluted 1:1000 into VBMM and incubated at 37 °C with shaking until exponential phase (OD_600_ 0.6). Cultures were then centrifuged, and the supernatants were sterile-filtered using 0.22 µm filters. Filtered supernatants were freshly used for quantification of 3OC_12_-HSL, C_4_-HSL, and AQs signal molecules by using the reporter strains PA14-R3 (78), C_4_-HSL-Rep (79), and AQ-Rep (80), respectively, as previously described (81). Briefly, 5 µL of supernatants were added to 195 µL of PA14-R3 (OD_600_ = 0.045), C_4_-HSL-Rep (OD_600_ = 0.045), and AQ-Rep (OD_600_ = 0.1) cultures, respectively, in 96-well, black, clear-bottomed microtiter plates. The resulting microtiter plates were incubated at 37°C. Light emission (RLUs) and cell density (OD_600_) were measured after 4 h (for 3OC_12_-HSL) or 6 h (for C_4_-HSL and AQs) of incubation using an automated luminometer-spectrophotometer plate reader Spark10M (Tecan). RLU values were normalized to OD_600_. Dedicated calibration curves were generated by growing each reporter strain in the presence of increasing concentrations of the corresponding synthetic signal molecule (PQS was used for the AQ-Rep strain), and these curves were used to calculate the concentration of the different QS signal molecules in each culture supernatant. Experiments were conducted in technical and biological triplicates.

## Statistical Analyses

Statistical analyses were performed using GraphPad Prism Software v.10.1.1 and were calculated as specified in the figure captions.

## Acknowledgments

This study was partially supported by a MUR grant to AB (PRIN 2022, contract 2022E57Z3K, CUP E53D23009850006, Funded by the European Union – NextGenerationEU”). Hyp-ACB infrastructure (grants n. GA116154C8A94E3D and SM12117A75F3D1B2 to F.C., also supported by Horizon H2020 MOSBRI project-Molecular Scale Biophysics Research Infrastructure- and member of IR2-TECH Infrastructure Rome Technopole) and Sapienza University of Rome (RM123188F4549144 to S.R.) are gratefully acknowledged.

